# TriPer, an optical probe tuned to the endoplasmic reticulum tracks H_2_O_2_ consumption by glutathione

**DOI:** 10.1101/088351

**Authors:** Eduardo Pinho Melo, Carlos Lopes, Peter Gollwitzer, Stephan Lortz, Sigurd Lenzen, Ilir Mehmeti, Clemens F. Kaminski, David Ron, Edward Avezov

## Abstract

The fate of H_2_O_2_ in the endoplasmic reticulum (ER) has been inferred indirectly from the activity of ER localized thiol oxidases and peroxiredoxins, *in vitro*, and the consequences of their genetic manipulation, *in vivo*. Here we report on the development of TriPer, a vital optical probe sensitive to changes in the concentration of H_2_O_2_ in the thiol-oxidizing environment of the ER. Consistent with the hypothesized contribution of oxidative protein folding to H_2_O_2_ production, ER-localized TriPer detected an increase in the luminal H_2_O_2_ signal upon induction of pro-insulin (a disulfide bonded protein of pancreatic β-cells), which was attenuated by the ectopic expression of catalase in the ER lumen. Interfering with glutathione production in the cytosol by buthionine sulfoximine (BSO) or enhancing its localized destruction by expression of the glutathione-degrading enzyme ChaC1 in lumen of the ER, enhanced further the luminal H_2_O_2_ signal and eroded β-cell viability. Tracking ER H_2_O_2_ in live cells points to an unanticipated role for glutathione in H_2_O_2_ turnover.

**Significance statement:** The presence of millimolar glutathione in the lumen of the endoplasmic reticulum has been difficult to understand purely in terms of modulation of protein-based disulphide bond formation in secreted proteins. Over the years hints have suggested that glutathione might have a role in reducing the heavy burden of hydrogen peroxide (H_2_O_2_) produced by the luminal enzymatic machinery for disulphide bond formation. However, limitations in existing in vivo H_2_O_2_ probes have rendered them all but useless in the thiol-oxidizing ER, precluding experimental follow-up of glutathione’s role ER H_2_O_2_ metabolism.

Here we report on the development and mechanistic characterization of an optical probe, TriPer that circumvents the limitations of previous sensors by retaining specific responsiveness to H_2_O_2_ in thiol-oxidizing environments. Application of this tool to the ER of an insulin-producing pancreatic b-cells model system revealed that ER glutathione antagonizes locally-produced H_2_O_2_ resulting from the oxidative folding of pro-insulin.

This study presents an interdisciplinary effort intersecting cell biology and chemistry: An original redox chemistry concept leading to development of a biological tool, broadly applicable for in vivo studies of H_2_O_2_ metabolism in the ER. More broadly, the concept developed here sets a precedent for applying a tri-cysteine relay system to discrimination between various oxidative reactants, in complex redox milieux.

## Introduction

The thiol redox environment of cells is compartmentalized, with disulfide bond formation confined to the lumen of the endoplasmic reticulum (ER) and mitochondrial inter-membrane space in eukaryotes and the peri-plasmic space in bacteria and largely excluded from the reducing cytosol (Sevier and Kaiser, 2008). Together, the tripeptide glutathione and its proteinaceous counterpart, thioredoxin, contribute to a chemical environment that maintains most cytosolic thiols in their reduced state. The enzymatic machinery for glutathione synthesis, turnover and reduction is localized to the cytosol, as is the thioredoxin/thioredoxin reductase couple (Toledano et al., 2013). However, unlike the thioredoxin/thioredoxin reductase system that is largely isolated from the endoplasmic reticulum, several lines of evidence suggest equilibration of glutathione pools between the cytosol and ER.

Isolated microsomes contain millimolar concentrations of glutathione (Bass et al., 2004); an estimate buttressed by kinetic measurements (Montero et al., 2013). Yeast genetics reveals that the kinetic defect in ER disulfide bond formation wrought by lack of an important luminal thiol oxidase, ERO1, can be ameliorated by attenuated glutathione synthesis in the cytosol (Cuozzo and Kaiser, 1999), whereas deregulated import of glutathione across the plasma membrane into the cytosol compromises oxidative protein folding in the yeast ER (Kumar et al., 2011). Import of reduced glutathione into the isolated rat liver microsomal fraction has been observed (Banhegyi et al., 1999) and in a functional counterpart to these experiments, excessive reduced glutathione on the cytosolic side of the plant cell ER membrane compromised disulfide formation (Lombardi et al., 2012). In mammalian cells experimental mis-localisation of the reduced glutathione-degrading enzyme ChaC1 to the ER depleted total cellular pools of glutathione (Tsunoda et al.,2014), arguing for transport of glutathione from its site of synthesis in the cytosol to the ER. Despite firm evidence for the existence of a pool of reduced glutathione in the ER, its functional role has remained obscure, as depleting ER glutathione in cultured fibroblasts affected neither disulfide bond formation nor their reductive re-shuffling (Tsunoda et al., 2014).

The ER is an important source of hydrogen peroxide production. This is partially explained by the activity of ERO1, which shuttles electrons from reduced thiols to molecular oxygen, converting the latter to hydrogen peroxide (Gross et al., 2006). Alternative ERO1-independent mechanisms for luminal hydrogen peroxide production also exist (Konno et al., 2015), yet the fate of this locally generated hydrogen peroxide is not entirely clear. Some is utilized for disulfide bond formation, a process that relies on the ER-localized peroxiredoxin 4 (PRDX4) (Tavender and Bulleid, 2010; Zito et al., 2010) and possibly other enzymes that function as peroxiredoxins (Nguyen et al., 2011; Ramming and Appenzeller-Herzog, 2013). However, under conditions of hydrogen peroxide hyper-production (experimentally induced by a deregulated mutation in ERO1), the peroxiredoxins that exploit the pool of reduced protein thiols in the ER lumen as electron donors, are unable to cope with the excess of hydrogen peroxide and cells expressing the hyperactive ERO1 are rendered hypersensitive to concomitant depletion of reduced glutathione (Hansen et al., 2012). Besides, ERO1 overexpression leads in increase of cell glutathione content (Molteni et al., 2004). These findings suggest a role for reduced glutathione in buffering excessive ER hydrogen peroxide production. Unfortunately, limitations in methods for measuring changes in the content of ER luminal hydrogen peroxide have frustrated efforts to pursue this hypothesis.

Here we describe the development of an optical method to track changes in hydrogen peroxide levels in the ER lumen. Its application to the study of cells in which the levels of hydrogen peroxide and glutathione were selectively manipulated in the ER and cytosol revealed an important role for glutathione in buffering the consequences of excessive ER hydrogen peroxide production. This process appears especially important to insulin-producing β-cells that are encumbered by a heavy burden of ER hydrogen peroxide production and a deficiency of the peroxide degrading calatase.

## Results

### Glutathione depletion exposes the hypersensitivity of pancreatic β-cells to hydrogen peroxide

Insulin producing pancreatic β-cells are relatively deficient in the hydrogen peroxide-degrading enzymes catalase and GPx1 (Lenzen et al., 1996; Tiedge et al., 1997) and thus deemed a sensitized experimental system to pursue the hypothesized role of glutathione in ER hydrogen peroxide metabolism. Compared with fibroblasts, insulin-producing RINm5F cells (a model for pancreatic β-cells) were noted to be hypersensitive to inhibition of glutathione biosynthesis by buthionine sulphoximine (BSO, Figure 1A & Figure S1). Cytosolic catalase expression reversed this hypersensitivity to BSO (Figure 1B & C).

**Figure 1.**
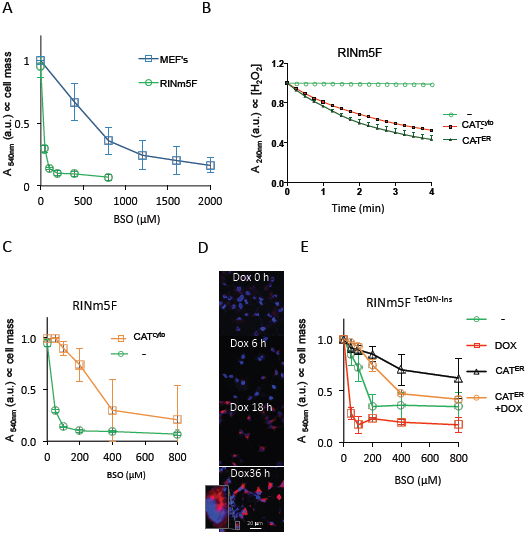
Glutathione depletion sensitizes pancreatic b-cells to endogenous H_2_O_2_.

(A) Absorbance at 540 nm (an indicator of cell mass) by cultures of a β-cell line (RINm5F) or fibroblasts (MEFs, a reference) that had been exposed to the indicated concentration of buthionine sulfoximine (BSO) before fixation and staining with crystal violet. (B) Plot of *in vitro* catalase activity, reflected in time-dependent decline in absorbance (A 240 nm) of H_2_O_2_ solution, exposed to lysates of untransfected RINm5F cells (-) or cells stably transfected with plasmids encoding cytoplasmic (CAT^cyto^) or ER-localized catalase (CAT^ER^). (C) As in (a), comparing un-transfected RINm5F cells (-) or cells stably expressing cytosolic catalase (CAT^cyto^). (D) Fluorescent photomicrographs of RINm5F cells stably expressing a tetracycline inducible human pro-insulin gene (RINm5F^TetON-Ins^) fixed at the indicated time points post doxycycline (20 ng/ml) exposure and immunostained for total insulin (red channel); Hoechst 33258 was used to visualize cell nuclei (blue channel). (E) As in (A) comparing cell mass of un-induced and doxycycline-induced RINm5F^TetON-Ins^ cells that had or had not been transfected with an expression plasmid encoding ER catalase (CAT^ER^). Shown are mean +/− SEM, n ≥ 3.

Induction of pro-insulin biosynthesis via a tetracycline inducible promoter (Figure 1D), which burdens the ER with disulfide bond formation and promotes the associated production of hydrogen peroxide, contributed to the injurious effects of BSO. But these were partially reversed by the presence of ER-localized catalase (Figure 1E). The protective effect of ER localized catalase is likely to reflect the enzymatic degradation of locally-produced hydrogen peroxide, as hydrogen peroxide is slow to equilibrate between the cytosol and ER (Konno et al., 2015). Together these findings hint at a role for glutathione in buffering the consequences of excessive production of hydrogen peroxide in the ER of pancreatic β-cells.

### A probe adapted to detect H_2_O_2_ in thiol-oxidizing environments

To further explore the role of glutathione in the metabolism of ER hydrogen peroxide we sought to measure the effects of manipulating glutathione availability on the changing levels of ER hydrogen peroxide. Exemplified by HyPer (Belousov et al., 2006), genetically encoded optical probes responsive to changing levels of hydrogen peroxide have been developed and via targeted localization, applied to the cytosol, peroxisome and mitochondrial matrix (Belousov et al., 2006; Gehrmann and Elsner, 2011; Gutscher et al., 2009; Malinouski et al., 2011). Unfortunately, in the thiol-oxidizing environment of the ER, the optically-sensitive disulfide in HyPer (that reports on the balance between hydrogen peroxide and contravening cellular reductive processes) instead forms via oxidized members of the protein disulfide isomerase family (PDIs), depleting the pool of reduced HyPer that can sense hydrogen peroxide (Konno et al., 2015; Mehmeti et al., 2012).

To circumvent this limitation we sought to develop a probe that would retain responsiveness to hydrogen peroxide in the presence of high concentration of oxidized PDI. HyPer consists of a circularly permuted yellow fluorescent protein (YFP) grafted with the hydrogen peroxide-sensing portion of the bacterial transcription factor OxyR (Belousov et al., 2006; Choi et al., 2001). It possesses two reactive cysteines: a peroxidatic cysteine (OxyR C199) that reacts with H_2_O_2_ to form a sulfenic acid and a resolving cysteine (OxyR C208) that attacks the sulfenic acid to form the optically distinct disulfide. We speculated that introduction of a third cysteine, vicinal to the resolving C208, might permit a rearrangement of the disulfide bonding pattern that could preserve a fraction of the peroxidatic cysteine in its reduced form and thereby preserve a measure of H_2_O_2_ responsiveness, even in the thiol-oxidizing environment of the ER.

Replacement of OxyR alanine 187 (located ~6 Å from the resolving cysteine 208 in PDB1I69) with cysteine gave rise to a tri-cysteine probe, TriPer, that retained responsiveness to H_2_O_2_ *in vitro* but with an optical readout that was profoundly different from that of HyPer: Whilst reduced HyPer exhibits a monotonic H_2_O_2_ and time-dependentincrease in its excitability at 488 nm compared to 405 nm (R^488/405^, Figure 2A), in response to H_2_O_2_, the R^488/405^ of reduced TriPer increased transiently before settling into a new steady state (Figure 2B). TriPer’s optical response to H_2_O_2_ was dependent on the peroxidatic cysteine (C199), as its replacement by serine eliminated all responsiveness (Figure 2C). R266 supports the peroxidatic properties of OxyR’s C199, likely by de-protonation of the reactive thiol (Choi et al., 2001). The R266A mutation similarly abolished H_2_O_2_ responsiveness of HyPer and TriPer indicating a shared catalytic mechanism for OxyR and the two derivative probes (Figure S2A).

**Figure 2.**
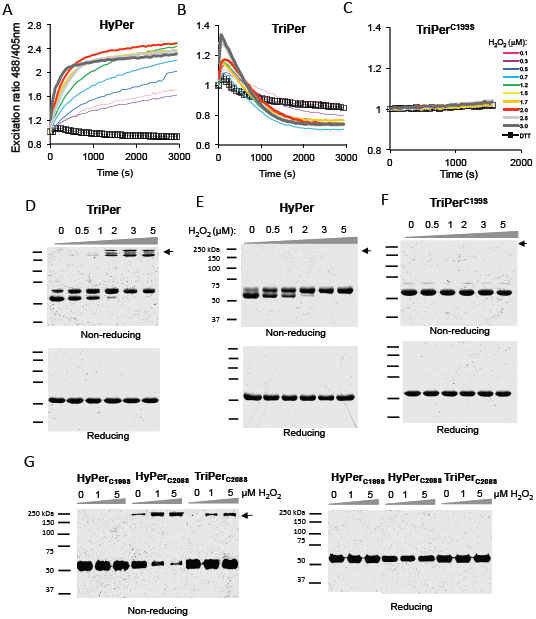
TriPer’s responsiveness to H_2_O_2_ *in vitro*.

(A)Traces of time-dependent changes to the redox-sensitive excitation ratio of HyPer, (B) TriPer or (C) the TriPer mutant lacking its peroxidatic cysteine (TriPer^C199S^), in response to increasing concentrations of H_2_O_2_ or the reducing agent dithiothreitol (DTT). (D) Non-reducing and reducing SDS-PAGE of recombinant TriPer, (E) HyPer, (F) a TriPer mutant lacking its peroxidatic cysteine (TriPer^C199S^) and (G) HyPer^C199S^ along HyPer/TriPer lacking their resolving cysteine (HyPer^C208S^/TriPer^C208S^) performed following the incubation *in vitro* with increasing concentrations of H_2_O_2_ for 15 min, black arrow denotes the high molecular weight species, exclusive to TriPer and to HyPer/TriPer lacking their resolving cysteine (HyPer^C208S^/TriPer^C208S^), emerging as a result of H_2_O_2_ induced dithiol(s) formation *in trans*. Shown are representatives of n ≥ 3.

The optical response to H_2_O_2_ of TriPer correlated in a dose and time-dependent manner with formation of high molecular weight disulfide bonded species, detectable on non-reducing SDS-PAGE (Figure 2D & Figure S3A). These species were not observed in H_2_O_2_-exposed HyPer and their presence in TriPer, depended on both the peroxidatic C199 and on R266 (Figure 2E, 2F and S2B). Furthermore, H_2_O_2_ promoted such mixed disulfides in probe variants missing the resolving C208 or both C208 and the TriPer-specific C187 (Figure 2G). The high molecular weight TriPer species induced by H_2_O_2_, migrate anomalously on standard SDS-PAGE, however on neutral pH gradient SDS-PAGE their size is consistent with that of a dimer (Figure S2C).

The observations above indicate that in the absence of C208, H_2_O_2_ induced C199 sulfenic intermediates are resolved *in trans* and suggest that formation of the divergent C208-C187 pair, unique to TriPer, favors this alternative route. To test this prediction we traced the R^488/405^ of TriPer’s under conditions mimicking oxidizing environment of the ER. TriPer’s time-dependent biphasic optical response (R^488/405^) to H_2_O_2_ contrasted with the hyperbolic profile of its response Diamide or PDI mediated oxidation (Figure 3A). The latter is by far the most abundant ER thiol-oxidizing enzyme. PDI catalyzed HyPer oxidation likewise had a hyperbolic profile but with a noticeable higher R^488/405^ plateau (Figure 3A). However, whereas TriPer retained responsiveness to H_2_O_2_, even from its PDI-oxidized plateau, PDI-oxidized HyPer lost all sensitivity to H_2_O_2_ (Figure 3B). Unlike H_2_O_2_, PDI did not promote formation of the disulfide bonded high molecular weight TriPer species (Figure S3A & S3B).

H_2_O_2_ driven formation of the optically active C199-C208 disulfide in HyPer enjoys a considerable kinetic advantage over its reduction by DTT (Konno et al., 2015). This was reflected here in the high R^488/405^ of the residual plateau of HyPer co-exposed to H_2_O_2_ and DTT (Figure 3C). Thus, HyPer and TriPer traces converge at a high ratio point in the presence of H_2_O_2_ and DTT (Figure 3C & S3C); convergence that requires both C199 and C208 (Figure S2D). In these conditions DTT releases TriPer’s C208 from the divergent disulfide, allowing it to resolve C199-sulfenic *in cis*, thus confirming C199-C208 as the only optically distinct (high R^488/405^) disulfide. It is worth noting that the convergence of TriPer and HyPer traces in these conditions confirms that in both C199-C208 in the sole high ratio states; consistent with the lack of optical response in all monomeric/dimeric configuration (Figure S2). Thus, TriPer’s biphasic response to H_2_O_2_, which is preserved in the face of PDI-driven oxidation (a mimic of conditions in the ER), emerges from the competing H_2_O_2_-driven formation of a *trans*-disulfide, imparting a low R^488/405^ (Figure 3D).

**Figure 3.**
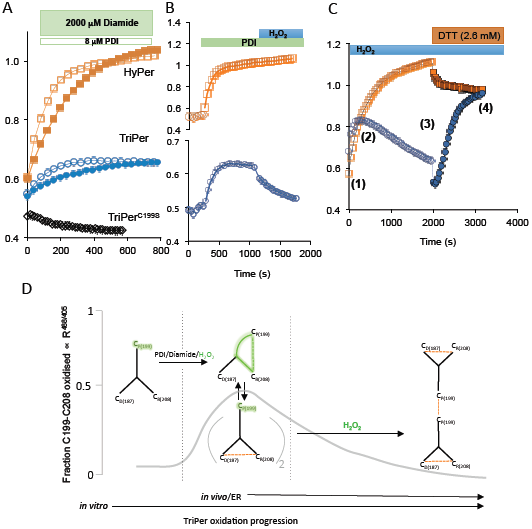
TriPer’s responsiveness to H_2_O_2_ in a thiol oxidizing environment.

(A)Traces of time-dependent changes to the excitation ratio of HyPer (orange squares), TriPer (blue spheres) and TriPer mutant lacking its peroxidatic cysteine (TriPer^C199S^, black diamonds), following the introduction of oxidised PDI (8 µM, empty squares, spheres and diamonds) or Diamide (general oxidant, 2 mM, filled squares and spheres). (B) Ratiometric traces (as in “a”) of HyPer and TriPer sequentially exposed to oxidized PDI (8 µM) and H_2_O_2_ (2 µM). (C) Ratiometric traces (as in “a”) of HyPer or TriPer exposed to H_2_O_2_ (4 µM) followed by DTT (2.6 mM). The excitation spectra of the phases of the reaction (1-4) are analyzed in Figure S3C. (D) Schema of TriPer oxidation pathway. Oxidants drive the formation of the optically distinct (high R^488/405^) C_P_199-C_R_208 (C_Peroxidatic_, C_Resolving_) disulfide which re-equilibrates with an optically-inert (low R^488/405^) C_D_187-C_R_208 (C_Divergent_, C_Resolving_) disulfide in a redox relay (Gutscher et al., 2008; Morgan et al., 2016; Sobotta et al., 2015) imposed by the three-cysteine system. The pool of TriPer with a reduced peroxidatic C_P_199 thus generated is available to react with H_2_O_2,_ forming a sulfenic intermediate. Resolution of this intermediate in *trans* shifts TriPer to a new, optically indistinct low R^488/405^ state depleting the optically distinct (high R^488/405^) intermolecular disulphide C_P_199-C_R_208. This accounts for biphasic response of TriPer to H_2_O_2_ (Figure 2B) and for its residual responsiveness to H_2_O_2_ after oxidation by PDI (Figure 3B).

### TriPer detects H_2_O_2_ in the oxidizing ER environment

To test if the promising features of TriPer observed *in vitro* enable H_2_O_2_ sensing in the ER, we tagged TriPer with a signal peptide and confirmed its ER localization in transfected cells (Figure. 4A). Unlike ER-localized HyPer, whose optical properties remained unchanged in cells exposed to H_2_O_2_, ER-localized TriPer responded with a H_2_O_2_ concentration-dependent decline in the R^488/405^ (Figure 4B & 4C).

The H_2_O_2_-mediated changes in the optical properties of ER-localized TriPer were readily reversed by washout or by introducing catalase into the culture media, which rapidly eliminated the H_2_O_2_ (Figure 4D & 4E). Both the slow rate of diffusion of H_2_O_2_ into the ER (Konno et al., 2015) and the inherent delay imposed by the two-step process entailed in TriPer’s responsiveness to H_2_O_2_ (Figure 3), contribute to the sluggish temporal profile of the changes observed in TriPer’s optical properties in cells exposed to H_2_O_2_. Further evidence that TriPer was indeed responding to changing H_2_O_2_ content of the ER was provided by the attenuated and delayed response to exogenous H_2_O_2_ observed in cells expressing an ER-localized catalase (Figure 4F).

**Figure.4.**
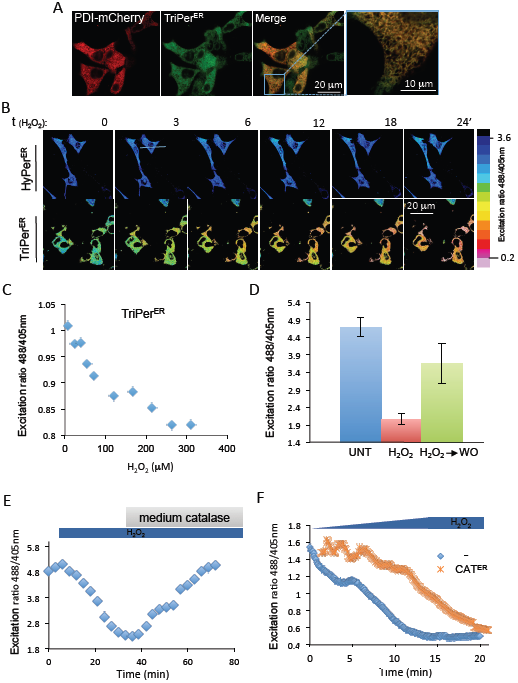
ER-localised TriPer responds optically to exogenous H_2_O_2._

(A) Fluorescent photomicrographs of live COS7 cells, co-expressing TriPer^ER^ and PDI-mCherry as an ER marker. (B) Time course of in vivo fluorescence excitation (488/405 nm) ratiometric images (R^488/405^) of HyPer^ER^ or TriPer^ER^ from cells exposed to H_2_O_2_ (0.2 mM) for the indicated time. The excitation ratio in the images was color coded according to the map shown. (C) Plot showing the dependence of in vivo fluorescence excitation (488/405 nm) ratio of TriPer^ER^ on H_2_O_2_ concentration in the culture medium. (D) Bar diagram of excitation ratio of TriPer^ER^ from untreated (UNT) cells, cells exposed to H_2_O_2_ (0.2 mM, 15 min) or 15 minutes after washout (H_2_O_2_∀WO) (Mean ± SEM, n=12). (E) A ratiometric trace of TriPer^ER^ expressed in RINm5F cells, exposed to H_2_O_2_ (0.2 mM) followed by bovine catalase for the indicated duration. (F) A ratiometric trace of TriPer^ER^ expressed alone or alongside ER catalase (CAT^ER^) in RINm5F cells, exposed to increasing concentrations of H_2_O_2_ (0 - 0.2 mM). Shown are representatives of n ≥ 3.

The response of TriPer to H_2_O_2_ could be tracked not only by following the changes in its excitation properties (as revealed in the R^488/405^) but also by monitoring the fluorophore’s fluorescence lifetime, using Fluorescent Lifetime Imaging Microscopy (FLIM) (as previously observed for other disulfide-based optical probes (Avezov et al., 2013; Bilan et al., 2013)).

Exposure of cells expressing ER TriPer to H_2_O_2_ resulted in highly reproducible increase in the fluorophore’s fluorescence lifetime (with a dynamic range > 8 X SD, Figure 5A). HyPer’s fluorescence lifetime was also responsive to H_2_O_2_, but only in the reducing environment of the cytoplasm (Figure 5B); the lifetime of ER localized HyPer remained unchanged in cells exposed to H_2_O_2_ (Figure 5C). These findings are consistent with nearly complete oxidation of the C199-C208 disulfide under basal conditions in ER-localized HyPer and highlight the residual H_2_O_2_-responsiveness of ER-localized TriPer (Figure 5D)(Konno et al., 2015; Mehmeti et al., 2012).

**Figure 5.**
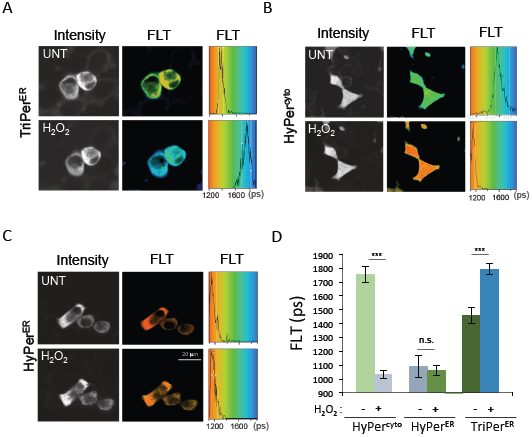
Fluorescence lifetime reports on the redox status of HyPer and TriPer

(A) Fluorescence intensity–based (in grayscale) and lifetime images (color-coded to the scale of the histogram on the right) of RINm5F cells expressing TriPer^ER^, (B) HyPer^cyto^ or (C) HyPer^ER^ before and after exposure to H_2_O_2_ (0.2 mM, 15 min). A histogram of the distribution of lifetimes in the population of cells is provided (right). (D) Bar diagram of the fluorescence lifetime peak values from (“A-C”, mean ± SD, ***P<0.005, n≥10).

Both ratiometry and FLIM trace alterations in the fluorophore resulting from C199-C208 formation. However, FLIM has important advantages over ratiometric measurements of changes in probe excitation, especially when applied to cell imaging: It is a photophysical property of the probe that is relatively independent of the ascertainment platform and indifferent to photobleaching. Therefore, though ratiometric imaging is practical for short-term tracking of single cells, FLIM is preferable when populations of cells exposed to divergent conditions are compared. Under basal conditions ER-localized TriPer’s lifetime indicated that it is found in a redox state where C199-C208 pair is nearly half-oxidized (Figure 5D), resembling that of PDI exposed TriPer in vitro (Figure 3A) and validating the use of FLIM to trace TriPer’s response to H_2_O_2_ in vivo.

### Glutathione depletion leads to H_2_O_2_ elevation in the ER of pancreatic cells

Exploiting the responsiveness of ER-localized TriPer to H_2_O_2_ we set out to measure the effect of glutathione depletion on the ER H_2_O_2_ signal as reflected in differences in ER TriPer’s fluorescence lifetime. BSO treatment of RINm5F cells increased the fluorescence lifetime of ER TriPer from 1490 +/− 43 at steady state to 1673 +/− 64 ps (Fig. 6A). A corresponding trend was also observed by measuring the changes in ER TriPer’s excitation properties ratiometrically (Fig. S4A).

Induction of pro-insulin biosynthesis accentuated TriPer’s response to BSO (Fig. 6A, compare samples 5 & 7), whereas expression of ER catalase counteracted the BSO induced increase of TriPer’s fluorescence lifetime, both under baseline conditions and following stimulation of pro-insulin production in RINm5FT^tetON-Ins^ cells (Fig. 6A compare samples 5 & 6 and 7 & 8, Fig 6B, compare samples 2 & 3). These observations correlate well with the cytoprotective effect of ER catalase in RINm5F cells exposed to BSO (Fig. 1). It is noteworthy that the increase in ER H_2_O_2_ signal in BSO-treated cells was observed well before the increase in the cytosolic H_2_O_2_ signal (Fig. 6C) and also preceded death of the glutathione depleted cells (Fig. S4B).

The ability of ER catalase to attenuate the optical response of ER-localized TriPer to BSO or pro-insulin induction argues for an increase in ER H_2_O_2_ as the underlying event triggering the optical response. Two further findings support this conclusion: 1) The disulfide state of the ER tuned redox reporter, ERroGFPiE (Avezov et al., 2013; Birk et al., 2013; van Lith et al., 2011) remained unaffected by BSO. This argues against the possibility that the observed TriPer response is a consequence of a more reducing ER thiol redox poise induced by glutathione depletion (Figure 6D). 2) TriPer’s responsiveness to BSO and pro-insulin induction was strictly dependent on R266, a residue that does not engage in thiol redox directly, but is required for the peroxidatic activity of TriPer C199 (Fig 6E & S4C). In addition, the above H_2_O_2_ specificity controls of TriPer response exclude other possible artificial affects on the probe’s fluorophore, such as pH changes.

**Figure 6.**
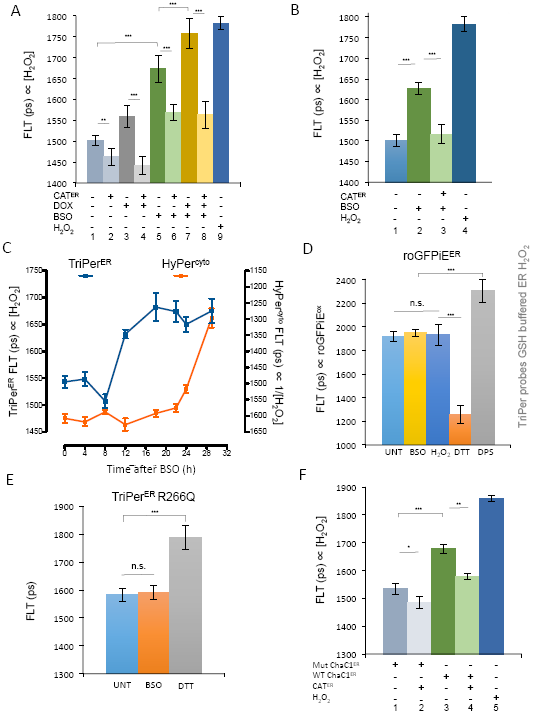
ER H_2_O_2_ increases in glutathione depleted cells.

(A) Bar diagram of fluorescence lifetime (FLT) of TriPer^ER^ expressed in the presence or absence of ER catalase (CAT^ER^) in RINm5F cells containing a tetracycline inducible pro-insulin gene. Where indicated pro-insulin expression was induced by doxycycline (DOX 20 ng/ml) and the cells were exposed to 0.3 mM BSO (18 hours) (B) As in “a” but TriPer^ER^-expressing cells were exposed to 0.15 mM BSO (18 hours. (C) A trace of time-dependent changes in HyPer^cyto^ or TriPer^ER^ FLT in RINm5F cells after exposure to 0.2 mM BSO. Each data point represents the mean ± SD of fluorescence lifetime measured in ≥ 20 cells. The ordinate of HyPer^cyto^ FLT was inverted to harmonize the trendlines of the two probes. (D) Bar diagram of FLT of ERroGFPiE, expressed in the presence or absence of ER catalase, untreated or exposed to 2 mM DTT, the oxidizing agent 2,2′-dipyridyl disulfide (DPS) or 0.3 mM BSO (18 hours). (E) Bar diagram of FLT of an H_2_O_2_ unresponsive TriPer mutant (TriPer^R266G^) expressed in the ER of untreated or BSO treated cells (0.3 mM; 18 hours). (F) Bar diagram of FLT of TriPer^ER^, expressed in the presence or absence of ER catalase or an ER localized glutathione-degrading enzyme (WT ChaC1^ER^) or its enzymatically inactive E116Q mutant version (Mut ChaC1^ER^). Shown are mean values ± SEM; *p<0.05, **p<0.01, ***p<0.005, *n* ≥ 20).

To further explore the links between glutathione depletion and accumulation of ER H_2_O_2_, we sought to measure the effects of selective depletion of the ER pool of glutathione on the ER H_2_O_2_ signal. ChaC1 is a mammalian enzyme that cleaves reduced glutathione into 5-oxoproline and cysteinyl-glycine (Kumar et al., 2012). We have adapted this normally cytosolic enzyme to function in the ER lumen and thereby deplete the ER pool of glutathione (Tsunoda et al., 2014). Enforced expression of ER localized ChaC1 in RINm5F cells led to an increase in fluorescence lifetime of ER TriPer, which was attenuated by concomitant expression of ER-localized catalase (Fig. 6F). Cysteinyl-glycine, the product of ChaC1 has a free thiol, but its ability to balance ER H_2_O_2_, may be affected by other factors such a clearance or protonation status. Given the relative selectivity of ER localized ChaC1 in depleting the luminal pool of glutathione (which equilibrates relatively slowly with the cytosolic pool (Kumar et al., 2011; Tsunoda et al., 2014)), these observations further support a role for ER-localized glutathione in the elimination ofluminal H_2_O_2_.

### Analysis of the potential for uncatalyzed quenching of H_2_O_2_ by the ER pool of glutathione

Two molecules of reduced glutathione can reduce a single molecule of H_2_O_2_ yielding a glutathione disulfide and two molecules of water, equation (1):

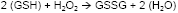

However there is no evidence that the ER is endowed with enzymes capable of catalyzing this thermodynamically-favored reaction. For while the ER possesses two glutathione peroxidases, GPx7 and GPx8, both lack key structural determinates for interacting with reduced glutathione and function instead as peroxiredoxins, ferrying electrons from reduced PDI to H_2_O_2_ (Nguyen et al., 2011). Therefore, we re-visited the feasibility of a role for the uncatalyzed reaction in H_2_O_2_ homeostasis in the ER.

Previous estimates of H_2_O_2_’s reactivity with reduced glutathione (based on measurements conducted in the presence of high concentrations of both reagents) yielded a rate constant of 22 M^-1^s^-1^ for the bi-molecular reaction (Winterbourn and Metodiewa, 1999). Exploiting the in vitro sensitivity of HyPer to H_2_O_2_ (Figure 2A) we re-visited this issue at physiologically-relevant conditions (pH 7.1, concentrations of reactants: ([GSH] <3 mM, [H_2_O_2_] < 10 µM) obtaining a similar value for the second order rate constant (29 ± 4 M^−1^s^−1^, Fig. 7A–7D).

**Figure 7.**
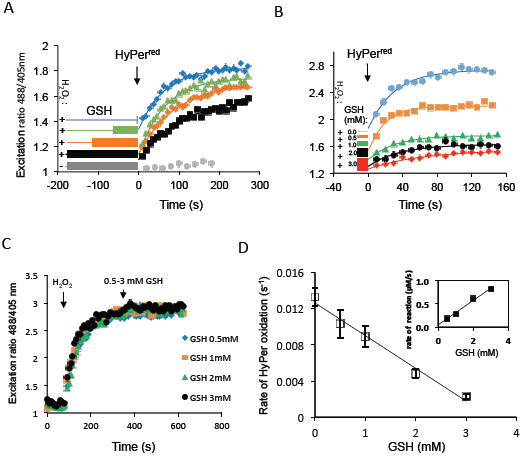
Kinetics of uncatalyzed elimination of H_2_O_2_ by reduced glutathione

(A) Traces of time-dependent changes to the excitation ratio (R^488/405^) of recombinant HyPer introduced into a solution of H_2_O_2_ (10 M) that had been preincubated with GSH (3mM) for increasing time periods, as indicated. Note the inverse relationship between HyPers oxidation rate and the length of the preceding H_2_O_2_-GSH pre-incubation. (B) As in (A) but following HyPer introduction into pre-mixed solutions of H_2_O_2_ (10 M) andreduced glutathione (GSH) at the indicated concentrations (0 – 3 mM). (C) As in (A) but HyPer was exposed sequentially to H_2_O_2_ (2 M) and various concentrations of GSH. (D) A plot of HyPer oxidation rate (reflecting the residual [H_2_O_2_]) as a function of GSH concentration. The slope (−0.0036 s^−1^ mM^−1^) and the Y-intercept (0.01263 s^−1^) of the curve were used to extract the dependence of H_2_O_2_ consumption rate (-d[H_2_O_2_]/dt, at 10µM H_2_O_2_) on each GSH concentration point (0-3 mM) according to equations (2) – (4). The bi-molecular rate constant was calculated by division of the slope of the resulting curve (shown in the inset) by the initial [H_2_O_2_] (10µM).

Considering a scenario whereby oxidative folding of pro-insulin (precursor of the major secretory product of β-cells) proceeds by the enzymatic transfer of two electrons from di-thiols to O_2_, (as in ERO1/PDI catalysis, generating one molecule of H_2_O_2_ perdisulfide formed (Gross et al., 2006)) and given, three disulfides in pro-insulin, a maximal production rate of 6*10^-4^ femtomoles pro-insulin/min/cell (Schuit et al., 1991) and an ER volume of 280 femtoliters (Dean, 1973), the resultant maximal generation rate of H_2_O_2_ has the potential to elevate its concentration by 0.098 µM per second. Given the rate constant of 29 M^−1^s^−1^ for the bi-molecular reaction of H_2_O_2_ with reduced glutathione and an estimated ER concentration of 15 mM GSH (Montero et al., 2013), at this production rate the concentration of H_2_O_2_ would stabilize at 0.23 µM, based solely on uncatalyzed reduction by GSH. Parallel processes that consume H_2_O_2_ to generate disulfide bonds would tend to push this concentration even lower (Nguyen et al., 2011; Ramming et al., 2014; Tavender and Bulleid, 2010; Zito et al., 2010), nonetheless this calculation indicates that GSH can play an important role in the uncatalyzed elimination of H_2_O_2_ from the ER.

## Discussion

The sensitivity of β-cells to glutathione depletion, the accentuation of this toxic effect by pro-insulin synthesis and the ability of ER catalase to counteract these challenges all hinted at a role for glutathione in coping with the burden of H_2_O_2_ produced in the ER. However, without means to track H_2_O_2_ in the ER of living cells, this would have remained an untested idea. TriPer has revealed that ER hydrogen peroxide levels increase with increased production of disulfide bonds in secreted proteins, providing direct evidence that the oxidative machinery of the ER does indeed produce H_2_O_2_ as a (bi)product. Less anticipated has been the increased H_2_O_2_ in the ER of glutathione-depleted cells. The contribution of glutathione to a reducing milieu in the nucleus and cytoplasm is well established, but it has not been easy to rationalize its presence in the oxidizing ER; especially as glutathione appears dispensable for the reductive step of disulfide isomerization in oxidative protein folding (Tsunoda et al., 2014). Consistently, the ER thiol redox poise resisted glutathione depletion. TriPer has thus pointed to a role for glutathione in buffering ER H_2_O_2_ production, providing a plausible benefit from the presence of a glutathione pool in the ER lumen.

TriPer’s ability to sense H_2_O_2_ in a thiol-oxidizing environment relies on the presence of an additional cysteine residue (A187C) near the resolving C208 in the OxyR segment. The presence of this additional cysteine attenuates the optical responsiveness of the probe to oxidation by PDI by creating a diversionary disulfide involving C208. Importantly, this diversionary disulfide, which forms at the expense of the (optically active) C199-C208 disulfide preserves a fraction of the peroxidatic C199 in its reduced form. Thus, even in the thiol-oxidizing environment of the ER a fraction of C199 thiolate is free to form a reversible H_2_O_2_ driven sulfenic intermediate that is resolved by disulfide formation *in trans*. Furthermore, by preserving a fraction of C199 in its reduced form, the diversionary C187-C208 disulfide also maintains a pool of C199 to resolve the C199 sulfenic to form the trans-disulfide bond. Thus, the H_2_O_2_ induced formation of the *trans* disulfide is a feature unique to TriPer, and it too is a consequence of the diversionary disulfide, which eliminates the strongly competing reaction of the resolving C208 *in cis* with the sulfenic acid at C199.

The ability of ER expressed TriPer to alter its optical properties in response to H_2_O_2_ is enabled through its semioxidized steady state, dictated by kinetically/quantitatively dominant PDI. In-vitro this stage can be reached by oxidants such as Diamide or PDI, while the second oxidation phase, with lowering R^488/405^, is exclusive to H_2_O_2._ The net result of the presence of a diversionary thiol at TriPer residue 187 (A187C) is to render the probe optically sensitive to H_2_O_2_ even in the presence of high concentrations of oxidized PDI, conditions in which the precursor probe, HyPer, loses all optical responsiveness.

The aforementioned theoretical arguments for TriPer’s direct responsiveness to H_2_O_2_ are further supported by empirical observations: the peroxidatic potential (unusual for cysteine thiols) of the probe’s C199 is enabled through a finally balance charge distribution in its vicinity, of which R266 is a crucial determinant (Choi et al., 2001). Eliminating this charge yielded a probe variant with its intact cysteine system, but unresponsive to H_2_O_2_. Further, the ability of ER catalase to reverse the changes in TriPer’s disposition argues that these are initiated by changes in H_2_O_2_ concentration.

Both Hyper and TriPer react with elements of the prevailing ER thiol redox buffering system (exemplified by their equilibration, in vitro, with PDI). In the case of HyPer this reactivity is ruinous, but even in the case of ER TriPer, which retains a modicum of sensitivity to H_2_O_2_, the elements of the complex kinetic regime that drive its redox state are not understood in quantitative terms. Thus, it is impossible to fully deconvolute the potential impact on TriPer of changes in the ER thiol redox milieu from changes in H_2_O_2_ concentration wrought by a given physiological perturbation - TriPer is sensitive to both. However, it is noteworthy that oxidation of TriPer by H_2_O_2_, leading to a mixed disulfide state, shifts the optical readout towards lower R^488/405^ and shorter fluorescent lifetimes. Whilst such shifts are also consistent with a surge in thiol reductive activity, it seems unlikely that exposure of cells to H_2_O_2_ results in a more thiol-reducing ER. Similar considerations apply to the state of the ER in cells depleted of glutathione, as it is hard to imagine how this would lead to a more thiol-reducing ER. Indeed the H_2_O_2_-insensitive redox probe roGFPiE, which is know to equilibrate with ER localized PDI, was unaffected by glutathione depletion. Thus, while we cannot formally exclude that glutathione depletion also affects TriPer’s redox status independently of changes in H_2_O_2_ concentration, the bulk of the evidence favors a role for TriPer in tracking the latter and in reporting on an increase in ER H_2_O_2_ in glutathione-depleted cells.

TriPer has been instrumental in flagging glutathione’s role in buffering ER luminal H_2_O_2_. This raises the question as to whether the thermodynamically favored reduction of H_2_O_2_ by GSH is accelerated by ER-localized enzymes or proceeds by uncatalyzed mass action. The cytoplasm and mitochondria possess peroxide consuming enzymes that are fueled by reduced glutathione (Lillig et al., 2008). However the ER lacks known counterparts. Such enzymes may be discovered in the future, as well as possible pathways of GSH mediated PDRX4 modulation. But meanwhile it is notable that the kinetic properties of the uncatalyzed reduction of H_2_O_2_ by GSH are consistent with its potential in keeping H_2_O_2_ concentration at bay.

An ER that eschews catalyzed reduction of H_2_O_2_ by GSH and relies instead on the slower uncatalyzed reaction may acquire certain advantages. At low concentrations of H_2_O_2_, an ER organized along these lines would be flexible to deploy a kinetically favored PRDX4-mediated peroxidation reaction to exploit H_2_O_2_ generated by ERO1 for further disulfide bond formation (Tavender et al., 2010; Zito et al., 2010). At higher concentrations of H_2_O_2_ PRDX4 inactivation (Wang et al., 2012) limits the utility of a PRDX4-based coping strategy (Tavender and Bulleid, 2010; Wang et al., 2012). However, the concentration-dependent reduction of H_2_O_2_ by GSH is poised to counteract a build up of ER H_2_O_2_. Whilst uncertainty the rate of H_2_O_2_ transported across the ER membrane (Appenzeller-Herzog et al., 2016), we favour a model whereby such transport is comparatively slow (Konno et al., 2015); which provides an explanation for the observed delay between the increase in ER and cytosolic H_2_O_2_ when glutathione synthesis was inhibited. Sequestration of H_2_O_2_ in the lumen of the ER protects the genome from this potentially harmful metabolite and enables the higher concentrations needed for the uncatalyzed reaction to progress at a reasonable pace.

The implementation of a probe that detects H_2_O_2_ in thiol oxidizing environments has revealed a remarkably simple mechanism to defend the cytosol and nucleus from a (bi)product of oxidative protein folding in the ER. This mechanism is especially important in secretory pancreatic β cells that are poorly equipped with catalase/peroxidase.

## Materials and Methods

### Plasmid construction

Table S1 lists the plasmids used, their lab names, description, published reference and a notation of their appearance in the figures.

**Table S1.** List of plasmids

| ID* | Plasmid name | Description | Reference | First appearance | Label in figure |
| --- | --- | --- | --- | --- | --- |
| 1820 | pLVX_TRE3G_hINS | Mammalian expression of tetracyclin inducible human proinsulin | This study | Figure 1 | Tet-Ins |
| 1824 | pLVX-EF1a-Tet3G | Regulator plasmid for Tet-On® 3G Inducible Expression System | Clontech | Figure 1 | Tet-Ins |
| 816 | HyPer_pSmt3_pET28b | Bacterial expression of HyPer | PMID: 26504166 | Figure 2 | HyPer |
| 1046 | TriPer_pSmt3_pet28b(MP2) | Bacterial expression of TriPer** | This study | Figure 2 | TriPer |
| 1045 | HyPerC199S_pSmt3_pET28b | Bacterial expression of HyPer lacking peroxidic cys, C199S | This study | Figure 2 | HyPerC199S |
| 1057 | TriPer_C199S_pSmt3_pet28b (MP2) | Bacterial expression of TriPer lacking peroxidic cys, C199S | This study | Figure 2 | TriPer C199S |
| 779 | HyPerC208S_pSmt3_pQE30 | Bacterial expression of HyPer lacking resolving cys, C208S | This study | Figure 2 | HyPer C208S |
| 1905 | TriPerC208S_pSmt3_pET28b | Bacterial expression of TriPer lacking resolving cys, C208S | This study | Figure 2 | TriPer C208S |
| 827 | HyPerR266A_pSmt3_pET28b | Bacterial expression of HyPer lacking peroxidic R266 | This study | Figure S2 | HyPer R266A |
| 1909 | TriPerR266A_pSmt3_pET28b | Bacterial expression of TriPer lacking peroxidic R266 | This study | Figure S2 | TriPer R266A |
| 233 | hPDI(18-508)pTrcHis-A | Bacterial expression of human PDI1A | PMID: 21145486 | Figure 3 | PDI |
| 887 | hPDI_WT_mCherry_KDEL-N3 (MP#1) | Mammalian expression of human PDI1A C-term fused to mCherry | PMID: 25575667 | Figure 4 | PDI-mCherry |
| 601 | pFLAG_ERHyPer_CMV1 | Mammalian expression of ER HyPer | PMID: 26504166 | Figure 4 | HyPerER |
| 1025 | pFLAG_ERTriPer_CMV1 | Mammalian expression of ER TriPer | This study | Figure 4 | TriPerER |
| 777 | pHyPer_cyto | Mammalian expression of cytoplasmic HyPer | PMID: 20692175 | Figure 5 | cytoHyPer |
| 1907 | pFLAG_ERTriPer_R266Q_CMV1 | Mammalian expression of ER TriPer lacking peroxidic R266 | This study | Figure S4 | TriPer R266Q |
| 527 | FLAGM1_roGFP_iE_pCDNA3 | Mammalian expression of ERroGFPiE | PMID: 23589496 | Figure 6 | ERroGFPiE |
| 988 | FLAGM1_mChac1_CtoS_mCherry_pCDNA5_FRT_TO | Mammalian expression of ER Chac1 C-term fused to mCherry | PMID: 25073928 | Figure 6 | Chac1 wt |
| 1028 | FLAGM1_mChac1_CtoS_E116Q_mCherry_pCDNA5 | Mammalian expression of inactive ER Chac1 C-term fused to mCherry | PMID: 25073928 | Figure 6 | Chac1 mut |

\*Unique plasmid identification number used internally (a lab number).

*Unique plasmid idenRficaRon number used internally (a lab number).
**Cysteine was introduced instead of alanine 187, which is found in vicinity fo C208 in the OxyR structure PDB1I69, PMID: 11301006; amino asid numbering as in OxyR

### Transfections, cell culture and cells survival analysis

Mouse embryonic fibroblasts (MEF), HEK293t and COS7 cells were cultured in DMEM; RINm5F cells (ATCC, Manassas, VA, USA) were cultured RPMI (Sigma, Gillingham, Dorset, UK), both supplemented with 10% fetal calf serum; and periodically confirmed to be mycoplasma free.

RINm5F cells containing a tetracycline/doxycycline inducible human pro-insulin gene (RINm5F^TetON-Ins^) were generated using Lenti-XTM Tet-On^®^ 3G system (Clontech, Saint-Germain-en-Laye, France), according to the manufacturers manual. RINm5F cells stably expressing cytoplasmic or ER-adapted(Lortz et al., 2015b) human catalase were described in (Konno et al., 2015; Lortz et al., 2015a).

Transfections were performed using the Neon Transfection System (Invitrogen, Paisley, UK) applying 3 µg of ERTriPer or ERHyPer DNA/1 x 10^6^ cells.

For the survival assays, 3 x 10^4^ cells per 35 mm well were plated and exposed to various concentrations of BSO starting 4 hours after the seeding for 48 hours (RINm5F cells) or 96 hours (MEF, HEK-293t, COS7 cells). At the end of this period BSO was washed out and the cells were allowed to recover up to the point where the untreated sample reached 90% confluence. Then the cells were fixed in 5% formaldehyde (Sigma, Gillingham, Dorset, UK) and stained with Gram’s crystal violet (Fluka-Sigma, Gillingham, Dorset, UK). For quantification of the stain incorporated into each sample (proportional to the cell mass) it was solubilized in methanol and subjected to absorbance measurements at 540 nm using Versamax microplate reader (Molecular Devices, Sunnyvale, CA, USA). The readouts of each set were normalized its the maximum value (untreated sample).

### Quantification of catalase enzymatic activity

Catalase enzyme activity was quantified as described earlier (Tiedge et al., 1998). Briefly, whole-cell extracts were homogenized in PBS through sonication on ice with a Braun-Sonic 125 sonifier (Braun, Melsungen, Germany). Subsequently the homogenates were centrifuged at 10,000 x g and 4 °C for 10 min. The protein content of the supernatant was assessed by the BC Assay (Thermo Fisher Scientific, Rockford, IL, USA). For quantification of the catalase enzyme activity 5 µg of the total protein lysate were added to 50 mmol/L potassium phosphate buffer (pH 7.8) containing 20 mmol/L H_2_O_2_. The specific catalase activity was measured by ultraviolet spectroscopy, monitoring the decomposition of H_2_O_2_ at 240 nm and calculated as described in (Tiedge et al., 1998).

### Immunofluorescence staining

Prior to immunofluorescence staining cells were fixed with 4% paraformaldehyde, permeabilized with 0.5% Triton X-100/PBS and blocked with 10% goat serum/PBS. Total mouse monoclonal anti-insulin IgG (I2018, clone K36AC10, Sigma, Gillingham, Dorset, UK) was used as primary and goat anti-rabbit IgG conjugated to DyLight 543 (Jackson ImmunoResearch Laboratories, West Grove, PA, USA) as secondary antibodies.

### Confocal microscopy, fluorescence lifetime imaging and image analysis

Cells transfected with the H_2_O_2_ reporters (HyPer or TriPer) were analyzed by laser scanning confocal microscopy system (LSM 780; Carl Zeiss, Jena, Germany) with a Plan-Apo-chromat 60x oil immersion lens (NA 1.4), coupled to a microscope incubator, maintaining standard tissue culture conditions (37°C, 5% CO_2_), in complete medium. Fluorescence ratiometric intensity images (512 ⨯ 512 points, 16 bit) of live cells were acquired. A diode 405 nm and Argon 488 nm lasers (6 and 4% output respectively) were used for excitation of the ratiometric probes in the multitrack mode with an HFT 488/405 beam splitter, the signal was detected with 506-568 nm band pass filters, the detector gain was arbitrary adjusted to yield an intensity ratio of the two channels to allow a stable baseline and detection of its redox related alterations.

FLIM experiments were performed on a modified version of a previously-described laser scanning multiparametric imaging system (Frank et al., 2007), coupled to a microscope incubator, maintaining standard tissue culture conditions (Okolab, Pozzuoli, Italy), using a pulsed (sub 10 ps, 20-40 MHz) supercontinuum (430-2000nm) light source (SC 450, Fianium Ltd., Southampton, UK). Desired excitation wavelength (typically 470nm) was tuned using an acousto-optic tunable filter (AA Opto-electronic AOTFnC-VIS). Desired emission was collected using 510/42 and detected by a fast photomultiplier tube (PMC-100, Becker & Hickl GmbH). Lifetimes for each pixel were recorded using time correlated single photon counting (TCSPC) circuitry (SPC-830, Becker & Hickl GmbH), keeping count rates below 1% of the laser repetition rate to prevent pulse pile-up. Images were acquired over 20 to 60s, with a typical flow rate of 5⨯10^4^ photons sec^−1^, avoiding the pile up effect. The data were processed using SPCImage (Becker & Hickl GmbH) fitting the time correlated photon count data obtained for each pixel of the image to a monoexponential decay function, yielding a value for lifetime on the pico-second scale.

After filtering out autofluorescence (by excluding pixels with a fluorescence lifetime that out of range of the probes) mean fluorescence lifetime of single cells was established. Each data point is constituted by the average and SD of measurements from at least 20 cells. Calculations of p. values were performed using two tailed t. test function in Microsoft Excel 2011 software.

### Protein purification and kinetic assays in vitro

For *in-vitro* assays, human PDI (PDIA1 18–508), HyPer and TriPer were expressed in the E. *coli* BL21 (DE3) strain, purified with Ni-NTA affinity chromatography and analyzed by fluorescence excitation ratiometry as previously described (Avezov et al., 2015). Briefly, HyPer, TriPer and their mutants were assayed *in vitro* in Tris-HCl buffer, pH 7.4, 150mM NaCl after being reduced with 50mM DTT for one hour followed by gel filtration to remove DTT.

To establish the reactivity of H_2_O_2_ with GSH different amounts of GSH (0-3 mM, pH adjusted to 7.1; Sigma, Gillingham, Dorset, UK) were mixed with 10 µM of H_2_O_2_ for a fixed time period, and then exposed to recombinant HyPer (2 µM, reduced by 40 mM DTT and gel filtered to remove DTT). The relationship between the rate of HyPer oxidation and [GSH], described by equation (2), was used to extract the remaining [H_2_O_2_] after its exposure to various [GSH] for a given period, using equation (3).

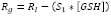

Where the rates of HyPer oxidation (s^−1^) in a given [H_2_O_2_] are denoted by R_g_ (GSH-affected) and R_i_ (in the absence of GSH), S_1_ is the experimental coefficient (s^−1^ mM^−1^). The latter two are the y-intercept and the slope of the curve in Figure 6D, accordingly.

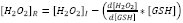

Where [H_2_O_2_]_R_ and [H_2_O_2_]_I_ are the residual and initial H_2_O_2_ concentrations accordingly. The resulting [H_2_O_2_] for each experimental [GSH] point were used to calculate the corresponding reaction rate according to equation (4), developed based on equation (3) for the special case of H_2_O_2_ – GSH reaction time (t) and [H_2_O_2_] (resulting curve shown in Fig. 6D inset, S_2_ is the slope). The bi-molecular rate constant (k) is given by equation (5).

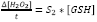

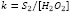

The concentration of H_2_O_2_ at the equilibrium where the rate of its supply equals the rate of its reaction with GSH was calculated according to the equation (6):

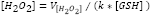

Where V_[H_2_O_2_]_ is the assumed rate of H_2_O_2_ generation and K is the bi-molecular rate constant for the GSH/H_2_O_2_ reactivity.

## Acknowledgments

Supported by grants from the Wellcome Trust (Wellcome 200848/Z/16/Z) and Fundação para a Ciência e Tecnologia, Portugal (PTDC/QUI/BIQ/119677/2010 and UID/BIM/04773/2013-CBMR) and, European Commission (EU FP7 Beta-Bat No: 277713) and, a Wellcome Trust Strategic Award for core facilities to the Cambridge Institute for Medical Research (Wellcome 100140). DR is a Wellcome Trust Principal Research Fellow.

---

**Fig. S1.**
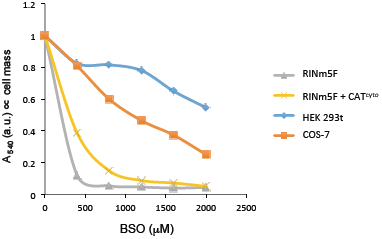
**Variable sensitivity of cultured cells to glutathione depletion.** As in Figure 1A, absorbance at 540 nm(an indicator of cell mass) by cultures of parental RINm5F, RINm5F stably over-expressing catalase in their cytosol (CAT^cyto^), HEK293 or COS7 cells that had been exposed to the indicated concentration of BSO before fixation and staining with crystal violet.

**Fig. S2.**
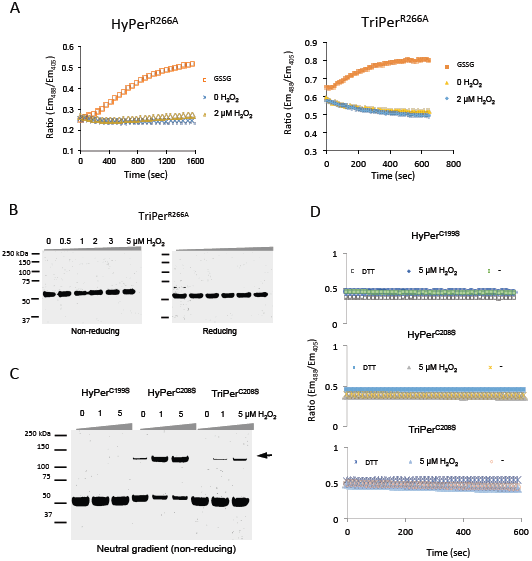
**The role of R266 in HyPerand TriPer’s reactivity with H_2_O_2_**. (A) Traces of time dependent changes to the excitation ratio of recombinant HyPer or TriPer variants with the inactivating R266A% mutation. (B) Non reducing and reducing SDS-PAGE of recombinant TriPer^R199A^ following incubation with increasing concentrations of H_2_O_2_ for 15 min. (C)Non-reducing gradient SDS-PAGE (pH 7.3, 4-12 %) of samples as in Figure 2G. (D)Traces of time-dependent changes to the excitation ratio of HyPer and TryPer mutant variants treated as in (B). Note, the variants lacking the ability to form C199-C208 disulfide do not change their excitation ratio upon oxidation.%

**Fig. S3.**
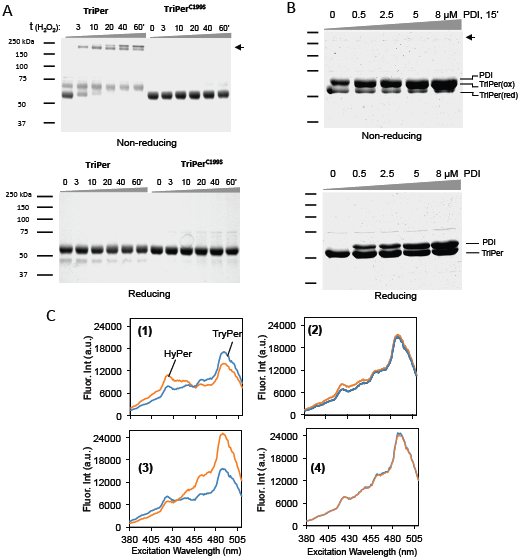
*In vitro*, H_2_O_2_ driven formation of disulphide-bonded high molecular TriPer species with divergent optical properties. (A) Coomassie-stained non-reducing and reducing SDS-PAGE of wildtype TriPer or its mutant variant lacking the peroxidatic cysteine (TriPer^C199S^) following exposure to 1.5µM of H_2_O_2_ for the indicated time period. Black arrows denote disulfide-bonded high molecular weight TriPer species. (B) As in “a” above, but following a 15 minute exposure to increasing concentrations of oxidized PDI (0C8%mM). Note the lack of the high molecular weight species in this sample and their prominance in the H_2_O_2_ treated sample (“a” above). (C) Excitation spectra (measured at emission 535 nm) of HyPer (orange trace) and TriPer (blue trace) for the different states of the probes, corresponding to phases 1-4 in Figure 3C.

**Fig. S4.**
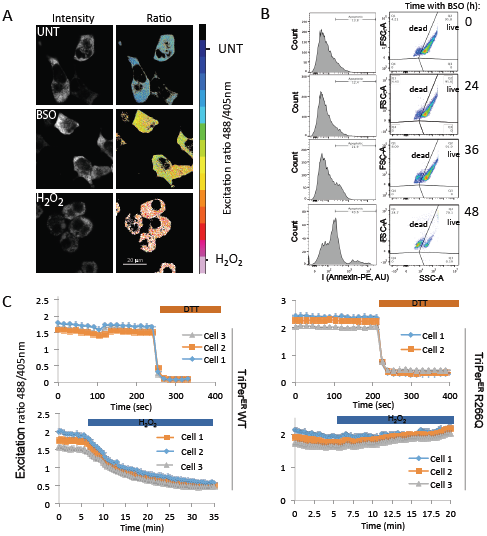
**Glutathione depletion induced apopto1c cell death and leads to concordant changes in TriPer^ER^ optical properties.** (A) Photomicrographs and fluoresce excitation% ratiometric images of untreated (UNT) RINm5F cellstransiently expressing TriPer^ER^ or cells treated with BSO (0.3 mM, 28 h) or H_2_O_2_ (0.2 mM, 15 min). The images were color coded for 488/405 nm excitation ratio (R^488/405^) according to the color map shown. (B) Flow cytometry analysis of RINm5F cells at the indicated Rme points aZer exposure to BSO (0.3 mM). Populations of dead and live cell were resolved by plonng forward vs. side scaoerings amplitudes (FCSCA and SSCCA accordingly). AppoptoRc cells populations were assessed by detecting surface phosphatidylserine using Phycoerythrin (PE) conjugated AnnexinV. Note that a significant population of dead cells only emerges aZer 36 hours, where as an increase in the ER H_2_O_2_ signal is observed by 12 hours (Figure 6C). (C) A ratiometric trace of TriPer^ER^ WT or TriPer^ER^ containing an R266Q mutation expressed in RINm5F cells, exposed to H_2_O_2_ (0.2 mM) or DTT (2mM) for the indicated duration.

